# Structural lipids enable the formation of functional oligomers of the eukaryotic purine symporter UapA

**DOI:** 10.1101/206714

**Authors:** Euan Pyle, Antreas C. Kalli, Sotiris Amillis, Zoe Hall, Aylin C. Hanyaloglu, George Diallinas, Bernadette Byrne, Argyris Politis

## Abstract

The role of membrane lipids in modulating eukaryotic transporter structure and function remains poorly understood. We used native mass spectrometry in combination with molecular dynamics simulations and *in vivo* analyses to investigate the roles of membrane lipids in the structure and transport activity of the purine transporter, UapA, from *Aspergillus nidulans*. We revealed that UapA exists mainly as a dimer and that two lipid molecules bind per UapA dimer. We identified three classes of phospholipids: phosphatidylcholine (PC), phosphatidylethanolamine (PE) and phosphatidylinositol (PI) which co-purified with UapA. Delipidation of UapA caused dissociation of the dimer into individual protomers. Subsequent addition of PI or PE rescued the UapA dimer and allowed recovery of bound lipids, suggesting a central role of these lipids in stabilising the dimer. We predicted a putative lipid-binding site near the UapA dimer interface. Mutational analyses established that lipid binding at this site is essential for formation of functional UapA dimers. Our findings reveal unprecedented level of detail into the nature of UapA-lipid interactions and provide a framework for studying similar eukaryotic systems.

## Introduction

The cellular membrane plays a key role in determining the structure and function of membrane proteins (Opekarova and Tanner 2003). Membranes are highly fluid and asymmetrical structures, incorporating discrete regions that provide distinct physical environments for the associated proteins (Engelman 2005). The biophysical properties of the membrane, including lateral and transverse pressures caused by membrane curvature and lipid packing, directly affect membrane protein folding, structure, and function (Marsh 1996; Booth and Curnow 2009; van den Brink-van der Laan et al. 2004). Furthermore, both specific and non-specific protein-lipid interactions can affect transporter conformation, stability, and oligomerisation (Laganowsky et al. 2014; Koshy and Ziegler 2015; Martens et al. 2016; Gupta et al. 2017). For example, specific lipid binding to subunit-subunit interfaces can modulate BetP oligomerisation (Koshy et al. 2013). In contrast, bulk or annular lipid interactions can provide structural support to facilitate conformational changes in the transporter NapA by stabilising the position of a static gate domain whilst allowing the movement of dynamic core domain to accommodate an elevator-like mechanism (Landreh et al. 2017). It is essential therefore to characterise protein-lipid interactions and the relationship between proteins and the cell membrane to fully understand the structure and function of membrane proteins *in vivo*.

Valuable insight has been gained by traditional structural biology methods, such as cryo-electron microscopy and X-ray crystallography; however, it is often difficult to capture protein-lipid interactions using these techniques (Peng, Yuan, and Hang 2014). Whilst there has been a marked increase in the number of membrane protein structures with resolved lipids in recent years, few studies have established the structure-function relationship of protein-lipid interactions (Mehmood et al. 2016; Norimatsu et al. 2017; Koshy et al. 2013). Recent studies have shown that native mass spectrometry (MS) can be used to detect, identify, and characterise protein-lipid interactions (Reading et al. 2015; Gupta et al. 2017; Laganowsky et al. 2013; Landreh et al. 2017; Mehmood et al. 2016; Skruzny et al. 2015; Zhou et al. 2011; Henrich et al. 2017). Native MS employs nano-electrospray ionisation (nESI) as a ‘soft’ ionisation technique which preserves the structure, oligomerisation, and ligand/lipid interactions of protein complexes (Hernandez and Robinson 2007; Ashcroft 2005). During native MS of intact membrane proteins, the detergent micelle is removed by collisional activation when protein ions are accelerated into a collision cell containing inert gas molecules. Careful tuning of the instrument parameters ensures well-resolved spectra, whilst maintaining the overall protein fold as well as interactions with other proteins, ligands and/or lipids. Consequently, native MS has emerged as a powerful tool for examining the role of lipid binding on the stability of membrane protein oligomers (Reading et al. 2015; Gupta et al. 2017).

Here, we used native MS to study UapA, a eukaryotic transporter from *Aspergillus nidulans* belonging to the nucleobase ascorbate transporter (NAT) family of metabolite importers. UapA is responsible for H^+^-dependent uptake of the purines xanthine and uric acid. The high-resolution structure of a transport inactive, conformationally locked mutant of UapA (G411V_Δ1-11_) was recently solved (Alguel et al. 2016) revealing that UapA is formed from two domains, the core domain and gate domain, and is likely to function via an elevator transport mechanism. The crystal structure also showed that UapA is a homodimer confirming earlier biochemical studies (Martzoukou et al. 2015), and, in combination with mutagenesis data, demonstrated that dimer formation was essential for function (Alguel et al. 2016).

Several recent studies using native MS have demonstrated that protein-lipid interactions play a crucial role in stabilising the dimer form of prokaryotic transporters (Gupta et al. 2017; Henrich et al. 2017). However, our understanding of the role of lipids in maintaining the quaternary structure of eukaryotic transporters remains limited. Here, we combined MS, molecular dynamics (MD), mutagenesis and functional analyses to establish that phosphatidylinositol (PI) and phosphatidylethanolamine (PE) are closely associated with UapA and play a crucial role in stabilising the functional UapA dimer.

## Results

### UapA expression, purification, and preliminary MS analysis

Wild-type (WT) UapA and a thermostabilised, conformationally-locked, inward-facing mutant, UapAG411V_Δ1-11_ (Leung et al. 2013), were individually expressed in *Saccharomyces cerevisiae* and purified in DDM to high homogeneity (**Supplementary Figure 1a**). Native MS experiments coupled to ion mobility (IM) (Michaelevski, Eisenstein, and Sharon 2010; Jurneczko and Barran 2011; Konijnenberg et al. 2014; Jenner et al. 2011; Ruotolo et al. 2008) report the overall molecular shape of biomolecules by determining their rotationally averaged collisional cross section (CCS). The CCS of UapAG411V_Δ1-11_ (6117Å) was found to be in agreement with the CCS of the folded, native-like state, calculated from the crystal structure of UapAG411V_Δ1-11_ (6230Å) (PDB: 5I6C) indicating that the protein remains folded in the gas phase (**Supplementary Figure 1b**).

### Protein-lipid interactions stabilise the UapA dimer

We performed native MS experiments on UapAG411V_Δ1-11_ revealing that the protein exists both as a monomer and a dimer (**Figure 1a**). The relative abundance of monomer:dimer was ^~^5:95 suggesting a strong dimer interface (Yefremova et al. 2017). The spectra also revealed the binding of two adduct molecules (measured mass: 775±50 Da) to the UapAG411V_Δ1-11_. We attributed this to the presence of bound phospholipids; however, the resolution of the peaks was insufficient to allow identification of the lipid species. We therefore extracted the lipids from the purified protein and analysed the lipid extract by liquid chromatography MS (LC-MS) and LC-MS/MS experiments. We identified phosphatidylcholine (PC), PE, and PI as the major lipid classes co-purifying with UapAG411V_Δ1-11_ (**Figure 1b** and **Supplementary Figure 2 and Supplementary Table 1**). We further verified the presence of the lipids identified by LC-MS in the UapAG411V_Δ1-11_ sample by negative ion native MS (**Figure 1c**). PE, PC, and PI are all abundant species in the plasma membranes of both the expression host (*S. cerevisiae*) and the native host (*A. nidulans*). Consequently, their interactions with UapA are likely to be physiologically relevant (Birch et al. 1998).

**Figure 1:**
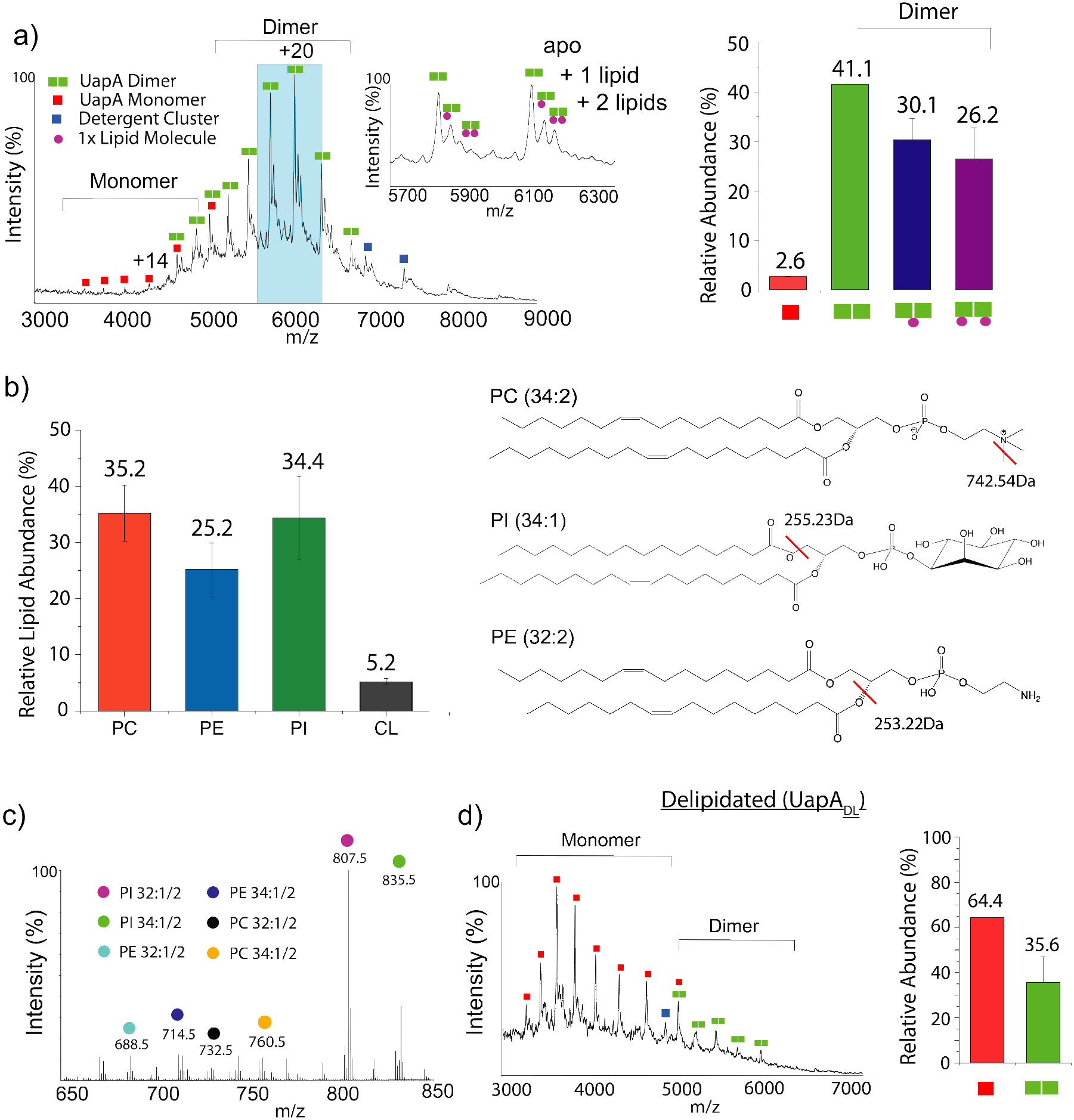
Bound lipids affect the oligomerisation of UapAG411V_Δ1-11_. (a) Mass spectrum of UapAG411V_Δ1-11_ highlighting the lipid binding peaks and the presence of both monomer and dimer in the gas phase. Inset shows a zoomed in view of the lipid binding peaks. The relative abundance of each UapA species is displayed in a bar chart (right). (b) Relative abundance of each lipid class identified by LC-MS and LC-MS/MS in a purified UapAG411V_Δ1-11_ sample and chemical structures of representative lipid species from each major class. Red lines indicate the fragmentation sites of each lipid identified by tandem MS (**Supplementary Figure 2**). (c) Negative ion mass spectrum of UapAG411V_Δ1-11_ in the low m/z range identifying the lipids co-purified with UapAG411V_Δ1-11_. Due to overlapping peaks of mono-unsaturated and poly-unsaturated fatty acid chains, both species are identified by 1/2 notation. (d) Mass spectrum showing the effect of delipidation on the relative abundance of UapAG411V_Δ1-11_ monomer and dimer. The relative abundance of each UapA species is displayed in a bar chart (right). The relative abundance of each oligomeric species in (a) and (d) was quantified using UniDec software (Marty et al. 2015). The mass spectra are representative of three independent experiments carried out under identical conditions. The relative abundance data is the average ± s.d; n = 3.

To investigate the effect of lipids in the formation of UapA dimers, we delipidated UapAG411V_Δ1-11_ and subjected it to native MS. Strikingly, our results showed that the protein is present primarily in the monomeric form (**Figure 1d**), suggesting that the lipids are important in maintaining the physiological dimer, as previously seen for prokaryotic transporters (Gupta et al, 2017). Loss of peaks in the ^~^750m/z range of the native MS spectra further confirmed almost total removal of lipid from the samples (**Supplementary Figure 3a**).

To explore the effect of the different lipids identified in lipidomics on UapA dimerisation, we titrated PI, PE or PC individually, or in combination, into delipidated UapAG411V_Δ1-11_ (**Figure 2**). Addition of either PE or PI restored the UapA dimer with both lipids displaying similar levels of dimer recovery, yielding 73.1% and 59.7% dimer, respectively (**Figure 2**). The addition of an equimolar mixture of PI and PE to delipidated UapAG411V_Δ1-11_ was more efficient at dimer recovery, yielding 81.5% dimer (**Supplementary Figure 3c**). This implies that the effects of the two lipids are additive. As a negative control we also titrated phosphatidylglycerol (PG), a lipid not usually found in eukaryotic plasma membranes, into delipidated UapAG411V_Δ1-11_ (Opekarova and Tanner 2003). PG did not induce any significant changes in the oligomerisation of delipidated UapAG411V_Δ1-11_ (**Supplementary Figure 3b**).

In contrast to PI and PE, the addition of PC to delipidated UapAG411V_Δ1-11_ failed to mediate the reformation of the UapA dimer. Indeed, adding PC actively induced dissociation of non-delipidated UapAG411V_Δ1-11_ (**Supplementary Figure 3d**). PC also generated an unknown UapAG411V_Δ1-11_ species (^~^69kDa) which we speculate to be a result of ^~^11 PC molecules binding non-specifically to the UapAG411V_Δ1-11_ monomer. Surprisingly, the presence of PC inhibited the dimer stabilising effects of PI and PE when mixtures of the three lipids were added to delipidated UapAG411V (**Supplementary Figure 3c**). Taken together, these data suggest that the dimer stabilising effects are specific to PI and PE lipids.

Next, we investigated the effects of lipid binding on the functionally active WT UapA. We found that the relative abundance of monomer:dimer was approximately 50:50 which is likely to be due to the reduced stability of WT UapA compared to the UapAG411V_Δ1-11_ (Leung et al. 2013) (**Supplementary Figure 4**). Delipidation of the wild-type protein resulted in almost complete loss of the dimer form and this could be partially recovered by addition of PI (**Supplementary Figure 4**).

**Figure 2:**
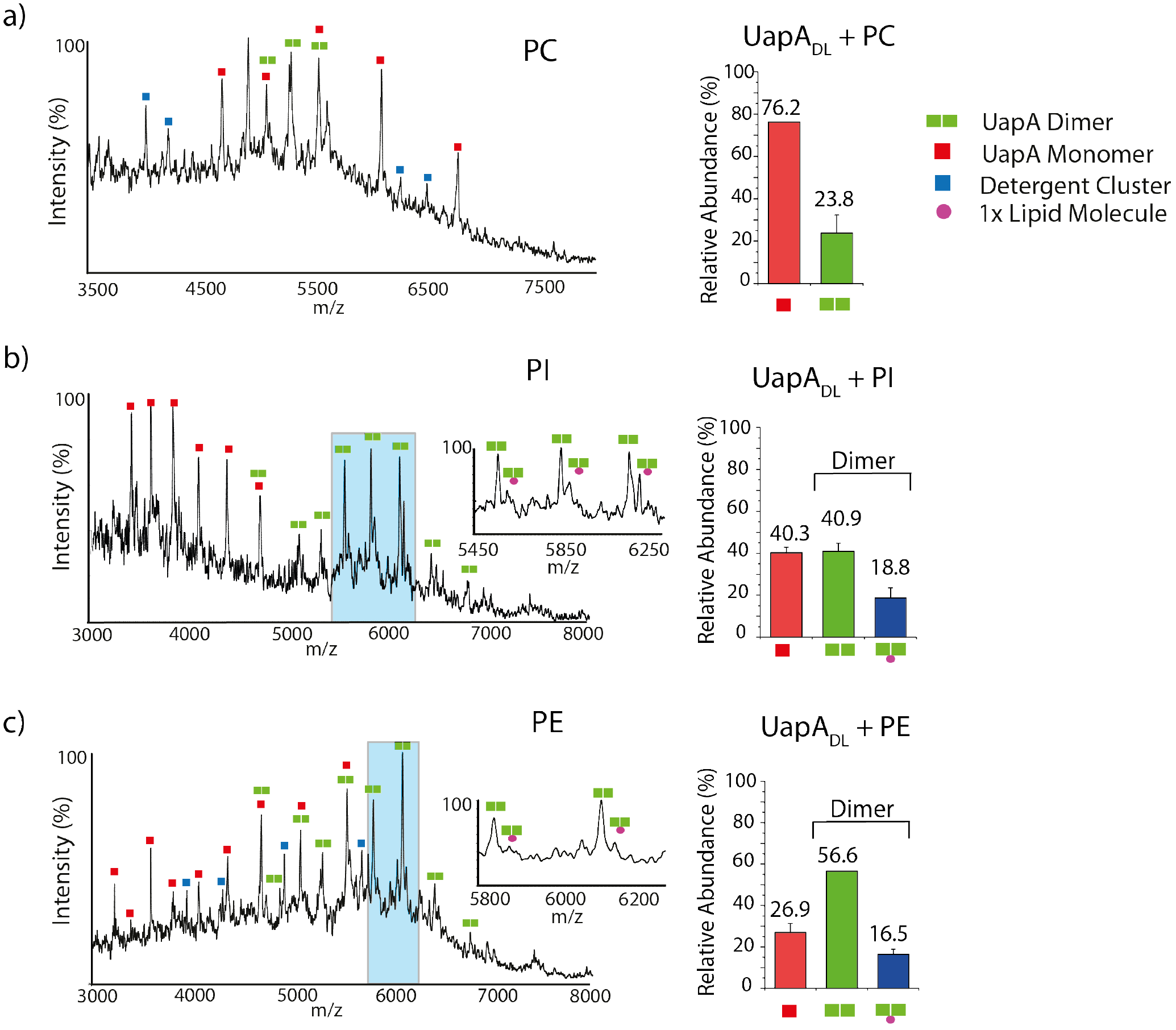
Mass spectra showing the effects on oligomerisation of adding (a) PC (34:1), (b) PI (34:1), and (c) PE (34:1) to delipidated UapAG411V_Δ1-11_. Lipid binding peaks are highlighted in the inset of the spectra. Lipids were added to delipidated UapAG411V_Δ1-11_ at a ratio of 100:1 lipid:UapA. The relative abundances of each oligomeric species were quantified using UniDec software (Marty et al. 2015). The mass spectra are representative of three independent experiments carried out under identical conditions. The relative abundance data are the average ± s.d; n = 3.

### MD simulations predict lipid binding sites at the dimer interface

MD simulations were performed in order to predict likely lipid binding sites. We carried out 5 μs simulations using a model of WT UapA based on the crystal structure of UapAG411V_Δ1-11_ (PDB: 5I6C). The structure was centred and embedded into a symmetric membrane containing PC (40%), PE (25%), and PI (35%) which closely resembles the lipid composition identified via lipidomics. Due to the long simulation time, we coarse-grained the molecules and used the MARTINI force field (Marrink and Tieleman 2013). In these simulations, the inward facing conformation of UapA was locked using an elastic network model (ENM). We analysed our simulations to predict any likely lipid binding sites near the dimer interface that may play a role in stabilising the dimer. Our MD simulations predicted that PC and PE interact with the same residues and that PI interacts with UapA residues with greater frequency than either PE or PC (**Supplementary Figure 5a**). No significant enrichment of PC or PE in the annual layer around UapA was observed. However, a number of positively charged Lys and Arg residues located principally on the intracellular side of the protein were predicted to be involved in PI binding (**Figure 3a**). Of particular interest was one clear PI binding site at the UapA dimer interface, comprised of residues Arg287 (at the cytoplasmic end of transmembrane (TM) 7, Arg478 and Arg479 (at the cytoplasmic end of TM13) and Asn539 (at the cytoplasmic end of TM14) (**Figure 3b**).

**Figure 3:**
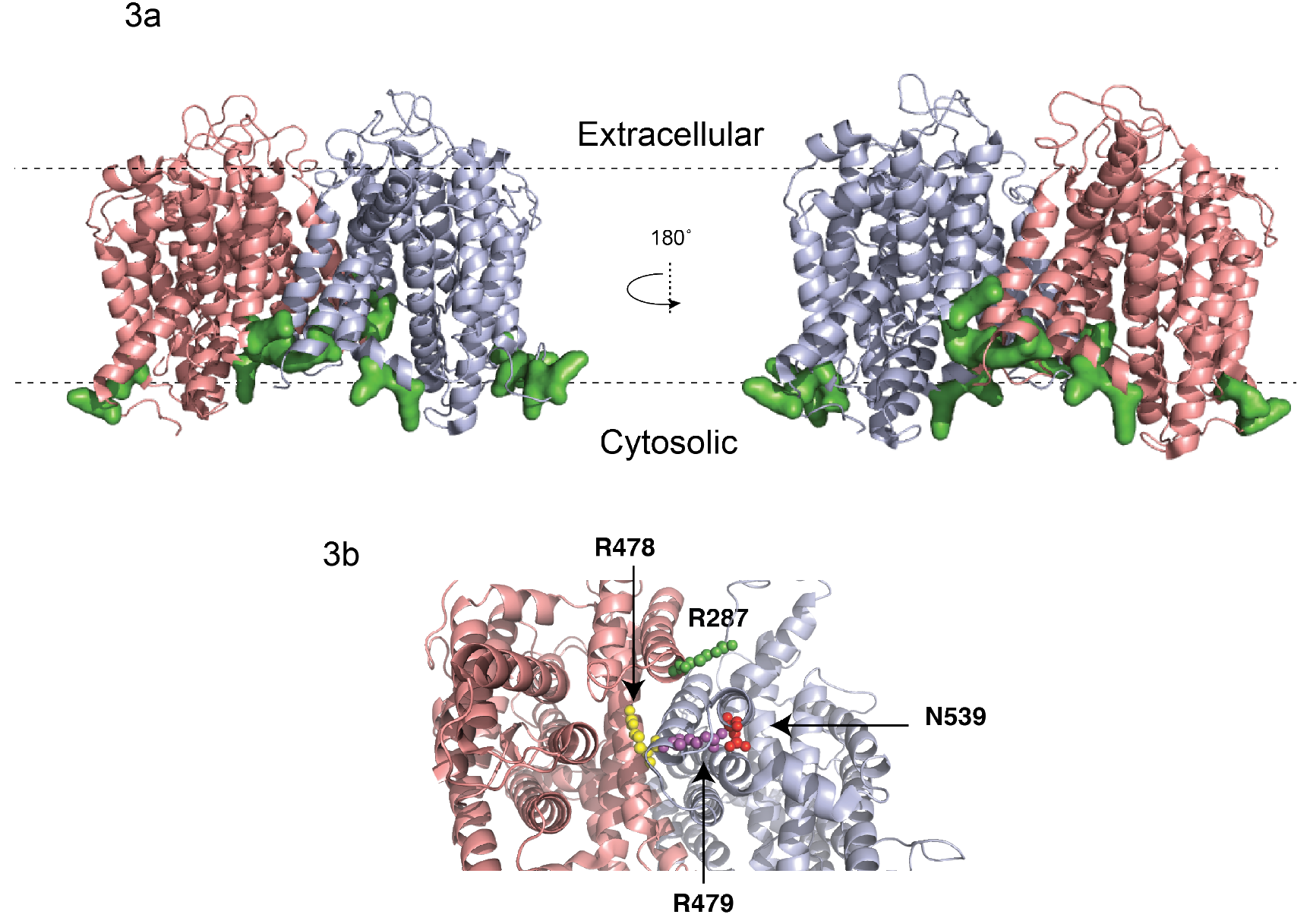
MD simulation of UapA in a symmetric bilayer consisting of PC:PE:PI 40:25:35. (a) Image of a cartoon representation of UapA with chain A coloured in red and chain B coloured in blue. Residues coloured in green had a normalised contact probability with PI higher than 0.8 over the course of the MD simulation (**Supplementary Figure 5b**), and were therefore predicted to have the highest number of interactions with the lipid. (b) Close-up of the putative lipid binding site near the dimer interface.

### Mutations to the putative lipid binding site cause loss of UapA function *in vivo*

In order to explore the role of the putative PI binding site in the structure and function of the UapA dimer we generated a range of single, double, and triple mutants of the residues (R287A, R478A, R479A, N539A, R478A/R479A, R287A/R478A/R479A) with C-terminal GFP tags. We individually transformed these constructs into a strain of *A. nidulans* (*uapAΔ uapCΔ azgAΔ pabaA1 argB2*) lacking all endogenous nucleobase transport systems. Transformants expressing the individual UapA mutants were tested for their capacity to grow using uric acid or xanthine as the sole nitrogen source. At 37°C, only the triple R287A/R478AR/R479A mutant showed markedly impaired growth on either uric acid or xanthine supplemented media, indicating a loss of UapA transport activity (**Figure 4a**). In contrast, at 25°C, closer to the physiological temperature of *A. nidulans*, both the triple R287A/R478AR/R479A and double R478A/R479A mutants were unable to grow on either uric acid or xanthine. Interestingly, the single R479A mutant also showed reduced growth on both substrates (**Figure 4a**).

Fluorescence microscopy carried out at 25°C showed that all of the mutants localized effectively to the plasma membrane confirming that loss of transport activity in the mutants is not a result of impaired UapA trafficking (**Figure 4b**). Further analysis using [^3^H] xanthine uptake assays (**Figure 4c and 4d**) confirmed the results from the growth assays and revealed close to wild-type substrate binding affinity for all mutants. This implies that the R478A/R479A and R287A/R478AR/R479A mutations affect substrate transport, rather than substrate binding. We have previously shown that dimer formation is critical for UapA function (Alguel et al. 2016), so in order to examine the oligomeric status of the UapA mutants, we carried out bimolecular fluorescence complementation (BiFC) analysis using a split YFP system (Martzoukou et al. 2015). This system uses two copies of the individual mutants co-expressed as fusions with either the N- or C-terminal domains of YFP. In this case, fluorescence is observed upon dimerization of copies of UapA tagged with the different YFP domains. Although the R287A/R478A/R479A mutant (UapA_3M_) was found to traffic to the membrane (**Figure 4b**), it was much less efficient at reconstituting the YFP (**Figure 4e**). This suggested that the UapA_3M_ mutant was principally trafficking as a monomer.

**Figure 4:**
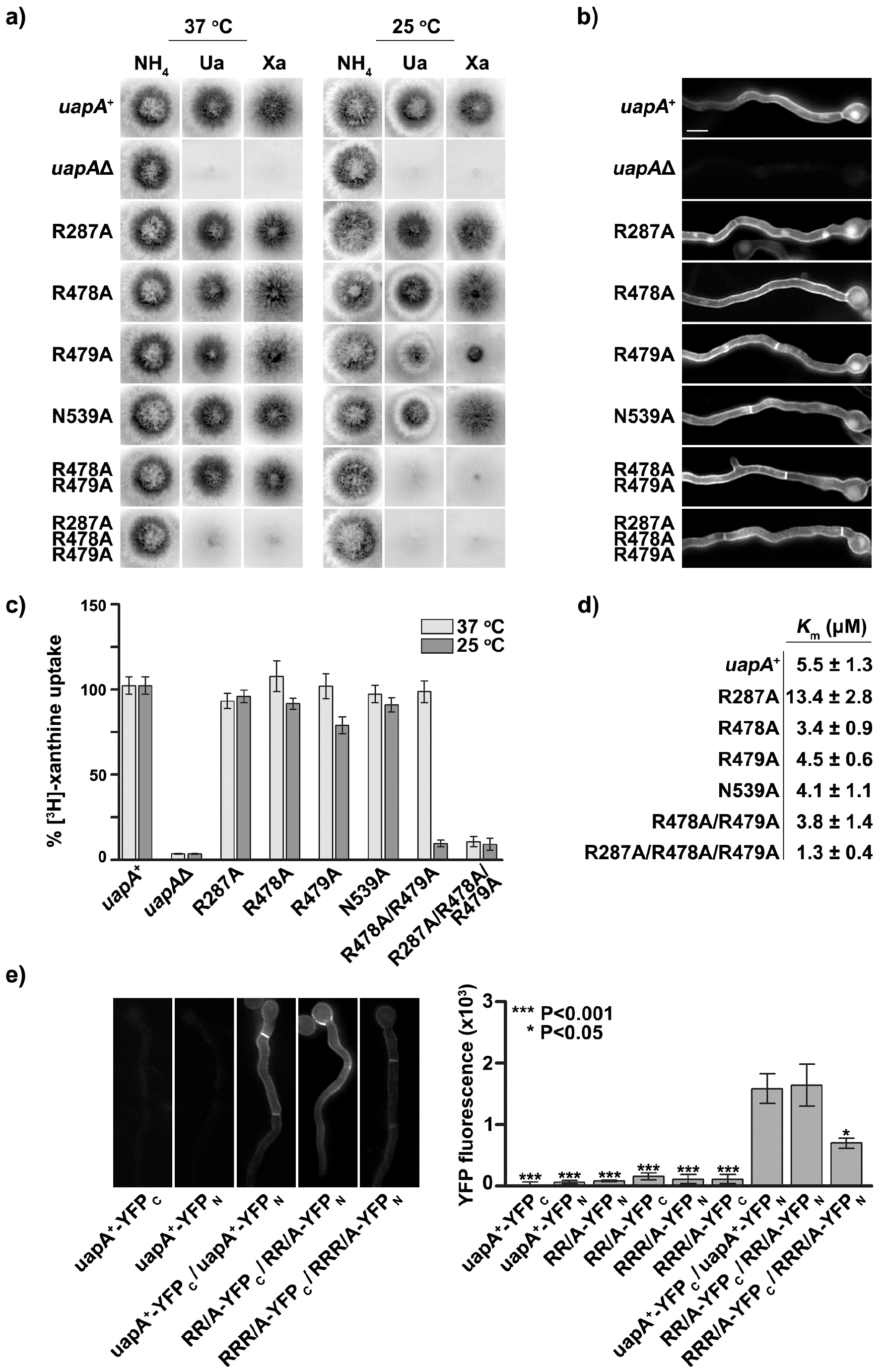
Effect of mutations to the putative lipid binding site on UapA function and trafficking in vivo. The constructs were transformed into the *A. nidulans* strain (*uapAΔ uapCΔ azgAΔ pabaA1 argB2*). uapAΔ indicates the original recipient strain as a negative control. (a) Growth tests of A. nidulans strains in minimal media supplemented with either ammonium (NH_4_^+^), uric acid (Ua), or xanthine (Xa) as the nitrogen source. Growth was assessed at 37°C and 25°C. (b) Inverted fluorescence microscopy images showing localization of the GFP tagged UapA constructs. All mutants display normal sorting to the membrane. (c) [^3^H]-Xanthine uptake assays, with rate of WT UapA uptake defined as 100%. d) Table of *K*_m_ values of the UapA mutants. (e) Bimolecular complementation (BiFC) analysis of the R478A/R479A (RR/A) UapA mutant and the R287A/R478A/R479A (RRR/A) UapA mutant. Mutant constructs tagged with the individual YFP domains were co-expressed in *A*. *nidulans*. Upon UapA dimerization the YFP is reconstituted. YFP fluorescence was measured by epifluorescence inverted microscopy. WT UapA (uapA+) expressed individually with either the C-terminal domain of YFP (YFPC) or the N-terminal domain of YFP (YFPN) are negative controls. Relative quantification of plasma membrane fluorescence intensity of mutants compared to WT UapA+−YFPC/UapA+−YFPN (right panel). The fungal growth and localisation data are representative of three independent experiments carried out under identical conditions. The *K*_m_ and fluorescence quantification date are the average ± s.d; n = 3.

In order to assess the precise oligomeric status of the UapA_3M_ mutant, we introduced the additional G411V mutation and 11 residue N-terminal truncation to generate the construct UapA_3M_ + G411V_Δ1-11_. Native MS of the purified UapA_3M_ + G411V_Δ1-11_ (**Supplementary Figure 1a**) showed a dramatic reduction in the abundance of UapA dimer compared to UapAG411V_Δ1-11_ (**Figure 5**). In addition, there was almost complete loss of lipid binding. This strongly suggests that the loss of UapA activity *in vivo* following disruption of the putative lipid binding site is due to the reduction of UapA-lipid interactions which play a key role in stabilising the functional dimer form.

**Figure 5:**
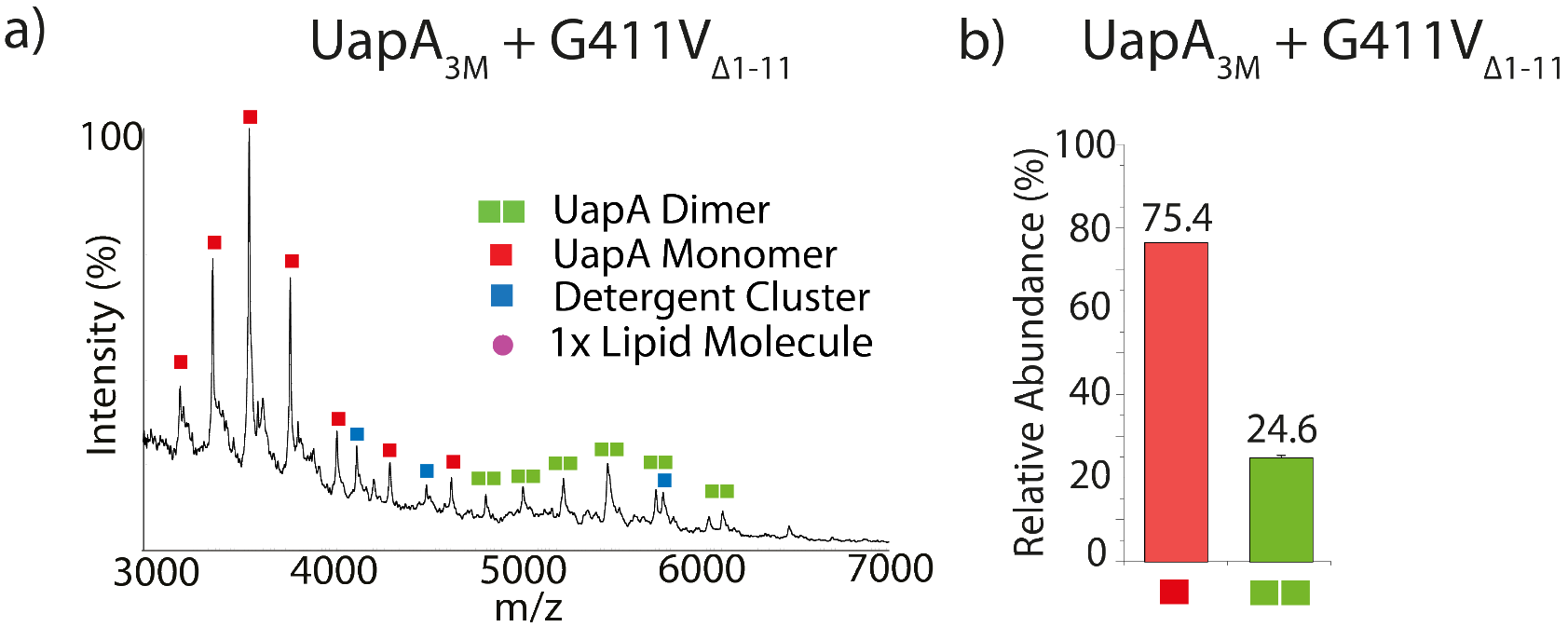
(a) Mass spectrum of UapA_3M_ with a G411V mutation and 11 residue N-terminal truncation. (b) Bar chart displaying the relative abundances of each oligomeric species of UapA_3M_ + G411V_Δ1-11_. See **Figure 1a** for the relative abundances of each oligomeric species of UapAG411V_Δ1-11_ under identical conditions in the mass spectrometer. The relative abundance of each species was quantified using UniDec software (Marty et al. 2015). The mass spectrum is representative of three independent experiments carried out under identical conditions. The relative abundance data is the average ± s.d; n = 3.

## Discussion

Recent years have seen a growing understanding of the key roles that membrane lipids have in the structure and function of membrane proteins; however, as yet there has been no in-depth study of the effects of lipids on eukaryotic transporters. Here, we explore the roles of lipids in maintaining the functional dimeric form of the eukaryotic UapA transporter. We combine native MS, lipidomics, MD simulations, mutagenesis and *in vivo* functional characterisation. Principally, we used the UapAG411V_Δ1-11_ construct, showing that it primarily exists as a dimer in the gas phase and that removal of tightly bound lipids causes dissociation into a monomer. The dimer can be recovered by addition of PI or PE. Importantly, we obtained similar results for WT UapA, although the spectra acquired for this protein are of reduced quality compared to the UapAG411V_Δ1-11_ construct due to the lower stability of the WT protein. However, this gave us confidence that further studies using the UapAG411V_Δ1-11_ were representative of the native protein.

In this study, protein-lipid interactions remained intact under high collision energies in the mass spectrometer. Therefore, it is unlikely that the lipid interactions we have described are annular, as these types of lipid usually bind in large numbers and are removed by low collisional energies (Landreh et al. 2016). Therefore, we speculate that PI and PE act as so-called structural lipids, forming integral interactions with UapA (Bechara and Robinson 2015). The dimer interface of UapA has a substantial surface area of ^~^6000 A^2^ (Alguel et al. 2016) which should render the dimer very stable; thus, the dissociation of the dimer into the monomeric form upon delipidation may be a result of the comparatively harsh treatment of the molecule rather than a direct effect of associated lipids. However, the fact that the dimer can be reformed by the addition of PI, PE or a combination of both lipids, is strongly suggestive that these lipids are having a specific stabilising effect on the oligomeric form of the transporter. There does not seem to be an absolute requirement for either PI or PE as individually both can recover dimer from delipidated protein to the same extent.

Interestingly, MD simulations predicted that most specific lipid binding sites are found within regions of the protein located on the intracellular side of the membrane and are therefore in contact with lipids in the inner membrane leaflet. Similar predictions have been recently made for NapA (Landreh et al. 2017) and CitS (Wöhlert et al. 2015) transporters. A high-resolution crystal structure of BetP also revealed asymmetry of lipids is important in mediating oligomer formation with a preference for lipids in the inner leaflet (Koshy et al, 2013). Of the four residues comprising the putative lipid binding site at the UapA dimer interface, only R479 is highly conserved among the eukaryotic NAT proteins. Interestingly, the R479A mutation was the only single substitution that had detectable effects on the *A. nidulans* growth rate. A loss of function observed for the R478A/R479A double mutant at the physiological temperature for *A. nidulans*, 25°C, and for the R287A/ R478A/R479A triple mutant at both 25°C and 37°C was not due to either a loss of correct trafficking to the membrane or impaired substrate binding. The apparent cryosensitivity of the single (R479A) and double (R478A/R479A) mutants may be due to a change in the *A. nidulans* lipid composition at the lower temperature. Such temperature dependent lipid composition changes have been reported for *S. cerevisiae* (Ejsing et al. 2009). Subsequent native MS analysis of the purified triple mutant in the G411V2_∆1-11_ background indicated that this construct is almost exclusively monomeric. Given that the functional form of UapA is the dimer, we reason that the lack of transport function in this protein is due to its an inability to bind lipids which promote dimerization.

It is important to note that both the native MS and the BiFC assay indicate that dimer does form in the absence of lipid binding, although to a much lower extent than when the lipid binding site at the dimer interface is intact. However, this form of the protein has lost virtually all transport activity. This indicates that lipid binding is not an absolute requirement for dimer formation but is essential for formation of the functional dimer.

The MD simulations predicted that lipids could also bind to the outermost, membrane facing, regions of the core domains of the UapA dimer. Lipid binding to this region has been predicted for NhaA and suggested to be involved in stabilising the core domain during the conformational transitions required for transport by the elevator mechanism (Landreh et al. 2017). However, in the case of UapA, removal of the putative lipid binding site at the dimer interface caused almost total loss of lipid binding as revealed by native MS (**Figure 5**). This indicates that binding to the core domain may not be a feature of UapA-lipid interactions. Further research is required to confirm this.

In conclusion, we have performed the first detailed analysis of the role of lipid binding to a eukaryotic transporter UapA, and shown that specific, structural lipids are critical for maintaining the protein in a functional, dimeric state. Overall, the combination of approaches used here have clear potential to allow investigation of the function of lipid binding to a range of different membrane proteins.

## Materials and methods

### Expression of the UapA constructs and membrane preparation

Wild-type (WT) UapA, a thermostable construct of UapA (UapAG411V_Δ1-11_), and a construct of UapA with disrupted lipid binding (R287A/R478A/R479A with G411V_Δ1-11_) (UapA_3M_) were recombinantly expressed as described previously (Alguel et al. 2016). In brief, *S. cerevisiae* UapA constructs were grown at 30°C and shaking at 300 rpm to FGY217 cells (6 or 12L) containing one of the an OD_600_ of 0.6. Expression was induced by the addition of galactose to a final concentration of 2%. After 22 hr incubation at 30°C with shaking the cells were harvested by centrifugation and resuspended in 10ml cell resuspension buffer (50 mM Tris, pH 7.6, 1 mM EDTA, 0.6 M sorbitol) per L culture. The cell suspension was flash frozen and stored at −80°C. Cells were lysed with a Constant Systems cell disruptor at 4°C. Unbroken cells and aggregates were removed by centrifugation at 4000 g for 10 minutes and membranes were isolated by ultracentrifugation at 100,000 g for 1 hr. Membranes were resuspended in membrane resuspension buffer (20mM Tris, pH 7.5, 0.3M sucrose, 0.1mM CaCl_2_) and flash frozen and stored at −80°C.

### Purification of UapA

Wild-type (WT) UapA, UapA_3M_, and UapAG411V_Δ1-11_ were purified as described previously (Alguel et al. 2016). In brief, membranes were solubilised for 1 hr at 4°C in membrane solubilisation buffer (1x PBS, pH 7.5, 10% glycerol (v/v), 1mM xanthine, 100mM NaCl, 1% n-dodecyl-β-D-maltoside (DDM_LA_) (v/v)) supplemented with 1 complete protease inhibitor tablet (Roche). Non-solubilised material was removed by centrifugation at 100,000 g for 45 min. The supernatant was incubated for 2 hr at 4°C with Ni^2+^-NTA superflow resin (Qiagen) equilibrated with Affinity buffer (1 x PBS, pH 7.5, 10% glycerol (v/v), 1 mM xanthine, 100 mM NaCl, 0.03% DDM_LA_ (v/v)) supplemented with 10 mM imidazole. The resin was packed onto a chromatography column and washed with 10 column volumes (CVs) Affinity buffer and then 30 CVs of Affinity buffer supplemented with 30 mM imidazole. UapA was eluted from the column with 5 CV Elution buffer (1x PBS, pH 7.5, 10% glycerol (v/v), 1 mM xanthine, 150 mM NaCl, 0.03% DDM_LA_ (v/v), 250 mM imidazole). Tobacco Etch Virus (TEV) protease was added to the protein at a protease:UapA ratio of 1:1, and the sample dialysed overnight into Dialysis buffer (20 mM Tris, pH 7.5, 5% glycerol (v/v), 0.6 mM xanthine, 150 mM NaCl, 0.03% DDM_LA_ (v/v)). The dialysed protein sample was loaded onto a 5 ml His-trap column (G.E. Biosciences) and UapA was recovered mainly in the initial flow through with any residual UapA eluted by washing the His-trap column with 5 CVs Reverse Affinity buffer (20 mM Tris, pH 7.5, 0.6 mM xanthine, 150 mM NaCl, 0.03% DDM_LA_ (v/v), 10 mM imidazole). The His-tagged GFP and His-tagged TEV remained bound to the column under these conditions. The UapA sample was then loaded onto a Superdex 200 10/300 gel filtration column pre-equilibrated with SEC buffer (20mM Tris, pH 7.5, 0.6mM xanthine, 150mM NaCl, 2x critical micellar concentration (CMC) of the detergent of interest). Fractions were analysed by SDS-PAGE. Fractions containing UapA were concentrated to ^~^9mg/ml. Protein solution was flash frozen and stored in small aliquots of 10μL at −80°C.

### Native MS

Capillaries for nESI were prepared using a Model P-97 (Sutter Instruments) capillary puller. Capillaries were gold coated using a Q150R S sputter coater (Quorum). Using Micro Bio-Spin 6 columns (Bio-Rad) purified UapA was buffer-exchanged into MS-compatible buffer (250 mM EDDA (pH 6.3), 0.014% DDM_LA_ (v/v), 10 mM L-serine) to a final protein concentration of 15 μM. The MS buffer was supplemented with 10 mM L-serine to improve spectral resolution by minimising salt adducts (Clarke and Campopiano 2015). Delipidated samples were prepared by incubating purified UapA with 1% DDM_LA_ (v/v) overnight before removing displaced lipids and buffer-exchanging UapA into the MS-compatible buffer using Micro Bio-Spin 6 columns (Bio-Rad). The individual proteins samples were loaded into gold-coated nanoflow capillaries (Hernandez and Robinson 2007) and introduced into a Synapt G2-Si (Waters) mass spectrometer by nESI. The conditions for maximum peak resolution for UapA were: capillary voltage +1.2-1.6 kV, sampling cone voltage 5 V, trap collision energy (CE) 125-200 V, transfer CE 50-240 V, backing pressure 3.88 mbar, trap and transfer pressure (argon) 1.72e^−2^ mbar, ion mobility cell pressure (nitrogen) 2.58mbar. A trap CE of 200 V and a transfer CE of 240 V was used when determining the relative abundances of each oligomeric species. To identify lipids in the protein sample, a negative capillary voltage of –1.1 kV was used. Mass measurements were calibrated using caesium iodide (100 mg/ml). Spectra were recorded and smoothed using Masslynx 4.1 (Waters) software.

### Native MS Lipid Titrations

Phosphatidylcholine (PC) (34:1), phosphatidylethanolamine (PE) (34:1), phosphatidylinositol (PI) (34:1), and phosphatidylglycerol (PG) (34:1) were supplied by Avanti Polar Lipids. Liposomes were prepared by dissolving lipid to 25 mg/ml in chloroform before evaporating the solvent under a N_2_ stream. The lipid film was frozen using liquid N_2_ and placed under vacuum in a freeze-dryer (Labconco Freezone) overnight. Lipids were solubilised to a concentration of 3 mM in 250 mM EDDA, 0.014% DDM_LA_ (v/v), 10 mM L-serine. Lipids were homogenised by 30 minutes sonication followed by three rounds of freeze-thawing. Lipids were added to delipidated UapA after buffer exchange into the MS-compatible buffer to a final concentration of 1.5mM lipid and 15μM UapA. Lipids were incubated with UapA for at least 1.5 hr before analysis by native MS under the following conditions: capillary voltage +1.2-1.6kV, sampling cone voltage 5V, trap CE 200V, transfer CE 240V, backing pressure 3.88 mbar, trap and transfer pressure (argon) 1.72e^−2^ mbar, ion mobility cell pressure (nitrogen) 2.58 mbar.

The relative abundances of each oligomeric state and lipid bound state of UapA was calculated using UniDec, a spectrum deconvolution software package (Marty et al. 2015), after correcting peak intensity by accounting for detector efficiency (Fraser 2002). Spectra were smoothed using MassLynx 4.1 (Waters) software prior to deconvolution. The mass range for peak detection was 61000-126000 Da. The following parameters were adjusted between samples to minimise the assignment of background noise as UapA species: charge (minimum of 8, maximum of 29-32), mass (monomer: 61025 ± 25, dimer 122025 ± 25, dimer + lipid 122775 ± 50, dimer + two lipids 123550 ± 50 Da), intensity threshold (0.0 to 0.45). This method assumes all UapA species have similar ionisation efficiencies.

### IM-MS

Conditions in the mass spectrometer for IM-MS of the monomer were: capillary voltage +1.2-1.6 kV, sampling cone voltage 5 V, trap CE 20 V, transfer CE 200 V, capillary backing pressure 0 bar. Drift times were measured at a T-wave height of 40 V and at three T-wave velocities (550, 600, 640 m/s). The following range of calibrants were utilised at 10 μM in 200 mM ammonium acetate: 10 μM concanavalin A, β-lactoglobulin, pyruvate kinase, glutamate dehydrogenase and alcohol dehydrogenase. PULSAR software (available from http://pulsar.chem.ox.ac.uk/) (Allison et al. 2015) was used to create a calibration curve (Bush et al. 2010) and calculate CCS values for UapA. MOBCAL software (available from http://www.indiana.edu/~nano/software/) was used to calculate the CCS of the UapA crystal structure (PDB: 5I6C) (Shvartsburg and Jarrold 1996). The CCS of missing residues from the crystal structure was accounted for using the following equation (Politis et al. 2010; Hall, Politis, and Robinson 2012):

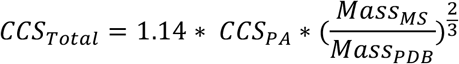

Equation 1: CCS_PA_ refers to the CCS calculated using the proximal approximation via MOBCAL. Mass_MS_ refers to the mass of the protein analysed by MS. Mass_PDB_ refers to the mass of protein in the PDB file.

### LC-MS

A 10μl 60μM UapA sample was subjected to a lipid extraction as described by the Folch method (Folch, Lees, and Sloane Stanley 1957). The lipid extract was separated by liquid chromatography using an Accela Autosampler (Thermo Scientific) coupled to a LTQ Orbitrap Elite (Thermo Scientific) mass spectrometer. 5μl lipid extract was separated at a flow rate of 0.5ml/min on an Acuity C18 BEH column (Waters, 50x2.1mm, 1.7μm) at 55°C. Mobile phase A = acetonitrile:water (60:40) with 10mM ammonium acetate. Mobile phase B = isopropanol:acetonitrile (90:10) with 10mM ammonium acetate. Lipids were initially separated with 60:40 mobile phase A:B. This mobile phase gradient was linearly changed to 1:99 A:B over 8 minutes and kept constant for 30 seconds. 60:40 mobile phase A:B was then regained in 10 seconds. These conditions were maintained for a further 2.5 minutes. Lipids were analysed by MS in the negative ion mode. Tandem MS was performed to fragment intact lipids, in order to identify the fatty acyl chains using their diagnostic ions and the LIPID MAPS database (Fahy et al. 2007). Cardiolipin also co-purified with UapAG411V_Δ1-11_ but is likely to be a contaminant from the yeast mitochondria (Schneiter et al. 1999).

### Coarse-grained Molecular Dynamics (CG-MD) Simulations

The CG-MD simulations were performed using the Martini 2.2 force-field (de Jong et al. 2013; Marrink et al. 2007) and GROMACS 4.6 (David Van Der et al. 2005). The crystal structure of UapA dimer (PDB: 5I6C, (Alguel et al. 2016); residues 66 to 545) was used for the CG-MD simulations. Missing unstructured regions from the crystal structure were added using Modeller (Fiser and Šali 2003; Sali and Blundell 1993) prior to the simulations and the V411 mutation in the crystal was mutated to glycine to generate a WT model. We also note that prior to the conversion to the coarse-grained representation, the substrate, xanthine, present in the crystal structure was removed.

A POPC bilayer was self-assembled around the UapA dimer and the snapshot at the end of this simulation was taken. Six different systems were generated with the protein inserted in complex symmetric bilayers that contained the following lipid concentrations: 1) 100% POPC, 2) 100% POPE, 3) 60% POPC-40% POPE, 4) 40% POPC-25% POPE-35% PI, 5) 65% POPC-25% POPE-10% PI, and 6) 70% POPC-25% POPE-5% PI. The exchange of lipids was done as previously described (Koldsø et al. 2014). For the simulations, we have used CG models of POPC and POPE lipids that had 4 CG particles in one of the lipids tails and 5 CG particles in the other tail (with one particle representing the double bond in the chain with 5 particles). Those are the CG-equivalent of the 1-palmitoyl 2-oleoyl- phosphatidylcholine (POPC) and 1-palmitoyl 2-oleoyl-phosphatidylethanolamine (POPE). The PI lipids had a tail with 4 CG particles and a tail with 5 CG particles but with no double bond. CG water particles were added to solvate the simulation systems and ^~^150 nM of NaCl was added to neutralize the systems. The systems were equilibrated for 10 ns with the protein backbone particles restrained. We ran simulations with both the dimer interface restrained with an ENM network and without restraining the dimer interface. For simulation systems 1 to 5 we have performed 5 repeat simulations in which an elastic network using a cut-off distance of 7 Å was used for the complete dimer and 5 repeat simulations in which the elastic network was applied to each UapA monomer. For simulation systems 6 and 7 we have performed simulations in which the elastic network was applied to each UapA monomer.

The integration step was 20 fs. All simulations were run for 5 μs each. The temperature was set to 323 K. The V-rescale thermostat (Bussi, Donadio, and Parrinello 2007) (coupling constant of 1.0) was used for temperature control. A Parrinello-Rahman barostat (Parrinello and Rahman 1981) (a coupling constant of 1.0 and a reference pressure of 1 bar) was used for pressure control. Lennard-Jones and Coulombic interactions were shifted to zero between 9 and 12 Å, and between 0 and 12 Å, respectively.

### *A. nidulans* Growth Conditions, UapA Localisation, Bifluorescence Complementation Assays and Xanthine Transport Assays

An *A. nidulans* mutant strain (*uapAΔ uapCΔ azgAΔ pabaA1 argB2*) lacking the genes encoding all major endogenous purine uptake systems was used (Pantazopoulou et al. 2007). This strain will only grow using xanthine or uric acid as the sole nitrogen source when a functional UapA construct is introduced. GFP-tagged UapA constructs (UapA^+^ (WT), R287A, R478A, R479A, N539A, R478A/R479A, R287A/R478A/R479A) were generated and transformed as previously described (Koukaki et al. 2003) into *A. nidulans*. The GFP tag has been shown not to affect UapA localisation, function, or transport kinetics in *A. nidulans* (Pantazopoulou et al. 2007). Successfully transformed strains of *A. nidulans* were selected via arginine auxotrophy complementation (Pantazopoulou et al. 2007).

As previously described (Karachaliou et al. 2013), UapA subcellular localisation was measured by visualising the GFP-tagged UapA using epifluorescence inverted microscopy (Zeiss Observer Z1/Axiocam HR R3 camera/Zen lite 2012 software). Radiolabelled [^3^H]-xanthine (22.8 Ci mmol^−1^, Moravek Biochemicals, CA, USA) uptake was measured using *A. nidulans* germinating conidiospores at 37°C or 25°C, pH 6.8, as previously described (Krypotou and Diallinas 2014). To measure growth of *A. nidulans* in different nitrogen sources, transformed strains were grown on minimal media supplemented with nitrogen sources (10mM ammonium tartrate or 0.5mM uric acid or 0.5mM xanthine) at 37°C or 25°C, pH 6.8. Dimerisation of mutant UapA was measured via a bimolecular fluorescence complementation assay, as described previously (Martzoukou et al. 2015). In brief, the N-terminal 154 amino acids of YFP or the C-terminal 86 amino acids of YFP were cloned into the pAN510exp or pAN520exp vector at the XbaI site. *uapA* with the necessary mutations was then cloned into the vector and transformed into *A. nidulans*. YFP fluorescence was measured using epifluorescence inverted microscopy (Zeiss Observer Z1/Axiocam HR R3 camera/Zen Lite 2012 software). Relative quantification using ICY and statistical analysis (Tukey’s Multiple Comparison Test, One-Way ANOVA for n=5 hyphae) of plasma membrane fluorescence intensity of mutants compared to WT UapA+−YFPC/UapA+−YFPN was performed as previously described (Martzoukou et al. 2017).

## Acknowledgements

The research was funded by the Wellcome Trust (109854/Z/15/Z) to AP and the Biotechnology and Biological Sciences Research Council grant BB/K017292/1 to BB. Work in the laboratory of GD was supported by a “Stavros S. Niarchos Foundation for Charity” grant. EP is the recipient of an Imperial College London Institute of Chemical Biology EPSRC CDT studentship awarded to BB, AP and ACH. We would like to thank Drs Eamonn Reading and Chloé Martens for their help in native MS experimental design and Dr Yilmaz Alguel for guidance in protein expression and purification. We thank Professor Julian Griffin for access to Orbitrap MS instrumentation. The MD simulations were undertaken on ARC2 and MARC1, part of the High Performance Computing and Leeds Institute for Data Analytics (LIDA) facilities at the University of Leeds, UK.

## Additional Information

### Abbreviations

BiFC, bimolecular fluorescence complementation; CCS, collisional cross section; CE, collision energy; CMC, critical micelle concentration; DDM, n-dodecyl-β-D-maltopyranoside; EDDA, ethylenediamine diacetate; IM-MS, ion-mobility mass spectrometry; LC-MS, liquid chromatography mass spectrometry; MD, molecular dynamics; MWCO, molecular weight cut off; NAT, nucleobase ascorbate transporter; nESI, nano-electrospray ionisation; OD_600_, optical density at 600nm; PC, phosphatidylcholine; PE, phosphatidylethanolamine; PG, phosphatidylglycerol; PI, phosphatidylinositol; UapA_3M_, UapA (R287A/R478A/R479A); WT, wild-type; YFP, yellow fluorescent protein.

### Author Contributions

The project was designed by A.C.H, B.B, and A.P.; E.P. expressed and purified the protein; Z.H. carried out lipidomics experiments; E.P. analysed the protein by native MS, and IM-MS; A.K. performed MD simulations; S.A. and G.D. carried out the mutagenesis functional analysis and localisation studies of the protein; E.P., B.B., and A.P. wrote the manuscript with contributions from all authors.

